# A2E induces the transactivation of RARs, PPARs and RXRs and its effects are counteracted by norbixin in retinal pigment epithelium cells *in vitro*

**DOI:** 10.1101/2020.03.30.016071

**Authors:** Valérie Fontaine, Mylène Fournié, Elodie Monteiro, Thinhinane Boumedine, Christine Balducci, Louis Guibout, Mathilde Latil, Pierre J. Dilda, José-Alain Sahel, Stanislas Veillet, René Lafont, Serge Camelo

**Affiliations:** Sorbonne Université, INSERM, CNRS, Institut de la Vision, 17 Rue Moreau, F-75012 Paris, France; Biophytis, Sorbonne Université, BC9, 4 place Jussieu, 75005 Paris, France; CHNO des Quinze-Vingts, DHU Sight Restore, INSERM-DGOS CIC 1423, 28 rue de Charenton, F-75012, Paris, France; Department of Ophthalmology, The University of Pittsburgh School of Medicine, Pittsburgh, PA, 15213, USA

**Author notes:** Corresponding author: Valérie Fontaine.

**Keywords:** N-retinylidene-N-retinylethanolamine (A2E), angiogenesis, apoptosis, inflammation, norbixin, nuclear receptor (NR), peroxisome proliferator-activated receptor (PPAR), retinoic acid receptor (RAR), retinal pigment epithelium (RPE), retinoic X receptor (RXR)

## Abstract

N-retinylidene-N-retinylethanolamine (A2E) plays a central role in age-related macular degeneration (AMD) by inducing apoptosis, angiogenesis and inflammation. It has been proposed that A2E effects are mediated at least partly via the retinoic acid receptor (RAR)-α. Here we show that A2E binds and transactivates not only RARs, but also peroxisome proliferator-activated receptors (PPARs) and retinoid X receptors (RXRs). Norbixin, which protects retinal pigment epithelium (RPE) cells against apoptosis induced by combined blue light illumination and A2E exposure, is also a ligand of these nuclear receptors (NRs) but does not induce their transactivation. Norbixin inhibits RXRs and PPARs but enhances RARs transactivation induced by A2E. Norbixin also inhibits PPAR-γ transactivation induced by its high affinity ligand troglitazone. Photoprotection of RPE cells by norbixin correlates with maintained levels of the antiapoptotic B-cell lymphoma 2 (Bcl2) protein. Moreover, norbixin reduces protein kinase B (AKT) phosphorylation, NF-κB and activator protein 1 (AP-1) transactivation, and the mRNA expression of the inflammatory interleukins (IL) 6 and 8 and of vascular endothelial growth factor (VEGF) that are enhanced by A2E. By contrast, norbixin increases matrix metalloproteinase 9 (MMP9) and C-C motif chemokine ligand 2 (CCL2) mRNA expression but has neither effect on extracellular signal-regulated kinase (ERK) phosphorylation, nor on IL-18 mRNA expression in response to A2E. Altogether, we show for the first time that A2E deleterious biological effects appear to be mediated through RARs, PPARs and RXRs. Moreover, we report that the modulation of these NRs by norbixin may open new avenues for the treatment of AMD.

## Introduction

AMD is the commonest cause of severe visual loss and blindness in developed countries among individuals aged 60 and older (1). A2E is a by-product of the visual cycle (2) formed by the reaction of 2 all-*trans* retinal molecules with phosphatidylethanolamine generating phosphatidylethanolamine-bisretinoid (A2-PE), as a detoxication mechanism of retinal isomers including all-*trans*- and 11-*cis*-retinal (3). A2E contained in RPE (4) and Bruch’s membrane (5) has been unambiguously associated with aetiology of AMD. *In vitro*, A2E in presence of blue light illumination is toxic for RPE (6, 7). Moreover, A2E alone increases the secretion of inflammatory cytokines by RPE cells *in vitro* but also in AMD animal models *in vivo* (8–10), and the expression of VEGF *in vitro* (11) and *in vivo* (12). However, knowledge on the mode of action of A2E explaining these effects is limited. It has been proposed that A2E is a ligand inducing the transactivation of the -α isoform of the RAR *in vitro* (11). In addition, it has been proposed that RAR-α is also responsible for VEGF production induced by A2E *in vitro* and *in vivo* (11, 12). Indeed, pharmacological inhibition of RARs transactivation with the RARs “specific” antagonist RO-41-5253 reduces A2E-induced VEGF production *in vitro* and *in vivo* and reduces the toxicity of A2E on RPE cells *in vivo* (12). However, it has been shown that RO-41-5253 is not only a RARs antagonist but also a partial agonist of PPAR-γ (13) suggesting that PPARs may also be involved.

The RARs isoforms α, β, and γ as well as the PPARs isoforms α, β/δ, and γ are NRs (14). In the presence of specific ligands, NRs become activated transcription factors which bind to specific DNA regulatory elements in the promoter or vicinity of target genes leading to their transcriptional activation or repression. Many NRs including RARs and PPARs form heterodimers with any of the three isoforms of the RXRs: RXR-α, -β or -γ (15, 16). Ligands of RARs alone are called retinoids and selective RXRs ligands are named rexinoids (16, 17). RARs/RXRs heterodimers are considered to be non-permissive because binding of an agonist to RARs subordinates the effect of agonist binding to RXRs (15, 16). Therefore, RARs/RXRs full transactivation by an RXRs agonist requires the concomitant binding of an agonist of RARs (18). Alternatively, RARs/RXRs heterodimers transactivation could be induced by a bi-specific agonist. By contrast, in the context of permissive heterodimers such as PPARs/RXRs or liver X receptors (LXRs)/RXRs, binding of an agonist of the specific NR partner or of an agonist of RXRs alone is sufficient to transactivate the NR partner (16, 19). RARs, RXRs and PPARs are known to regulate a wide spectrum of biological pathways ranging from metabolism, cell death, inflammation and angiogenesis (14) that are involved in AMD pathogenesis (20–22). Therefore, the role of NRs in A2E biological effects deserves additional studies.

9’-*cis*-Norbixin (norbixin) is a 6,6’-di-*apo*-carotenoid extracted from annatto (*Bixa orellana*) seeds (23). We have previously demonstrated that norbixin protects primary porcine RPE cells from phototoxicity induced by blue light illumination coupled with A2E-exposure *in vitro* (24). Moreover, norbixin reduces accumulation of A2E in RPE cells *in vitro* (24). Recently, it has been reported that norbixin is a PPAR-γ ligand with agonist activity (25). However, in another cellular context it has been shown that norbixin at 20 μM did not induce PPAR-γ transactivation in 3T3-L1 adipocytes (26). We wondered how norbixin could interfere with A2E toxic effects and if norbixin interactions with PPARs, RARs and RXRs were involved. To address these questions, we tested the effects of A2E in absence or presence of norbixin on RARs, PPARs and RXRs transactivation, and cell death, inflammatory molecules and VEGF expression mediated by A2E in primary porcine RPE cells *in vitro*. Here we show that A2E deleterious biological effects appear to be mediated through the transactivation of RARs, PPARs and RXRs. Moreover, we suggest that inhibition by norbixin of cell death, VEGF expression and inflammatory molecules induced by A2E is associated with modulation of the transactivation of these NRs.

## Results

### A2E binds RAR-α, RXR-α, PPAR-α and -γ and transactivates RARs, RXRs and PPARs

First, we wished to re-evaluate the binding profile of A2E on RAR-α, RXR-α, and PPARs. We determined whether A2E acts as a specific cognate ligand for RAR-α, RXR-α, PPAR-α, and PPAR-γ, by competition experiments *in vitro*. We confirmed previous observations by Iriyama and colleagues (11). We observed that A2E binds to RAR-α (Fig. 1A) with a Kd of 1.4 10^−6^ M and a half-maximal inhibitory concentration (IC50) of 2.79 10^−6^ M (Fig. 1B) whereas IC50 of all-*trans*-retinoic acid (ATRA), a pan agonist of RARs was 9.5 10^−9^ M in our hands (data not shown) and was previously reported to be 0.49 10^−9^ M [11]. The control vehicle alone did not interfere with the binding of [^3^H] ATRA to RAR-α, confirming the specific binding of A2E (data not shown). In addition, we demonstrated for the first time that A2E is also a ligand of RXR-α with an IC50 of 6.1 10^−6^ M whereas the IC50 of 9-*cis*-retinoic acid a high affinity RXR-α agonist is 94 10^−9^ M (data not shown). Interestingly, A2E binds also PPAR-α (Fig. 1C) and -γ (Fig. 1D) albeit with lower affinities (IC50 of 25.5 10^−6^ M, and 42.7 10^−6^ M respectively) than for RAR-α and RXR-α. Altogether the order of affinities of A2E for the NRs tested is as follow: RAR-α (Kd = 1.4 10^−6^ M) > RXR-α (Kd = 4.3 10^−6^ M) > PPAR-γ (Kd = 1.4 10^−5^ M) > PPAR-α (Kd = 4.1 10^−5^ M) and is reported in Table 1. Then the effects of A2E on the endogenous NRs transactivation in porcine RPE cells were investigated by means of a luciferase assay. According with its binding affinities, we observed in dose-response experiments that A2E transactivates RARs (Fig. 1E) and RXRs (Fig. 1F) at minimal concentrations of 5 and 20 μM respectively. The minimal concentration of A2E required to transactivate all PPARs in porcine RPE cells is 20 μM (Fig. 1G). Altogether our results show that A2E binds RAR-α and RXR-α and transactivates RARs and RXRs and that, despite its low affinity for PPARs, it is also able to induce their transactivation.

**Figure 1.**
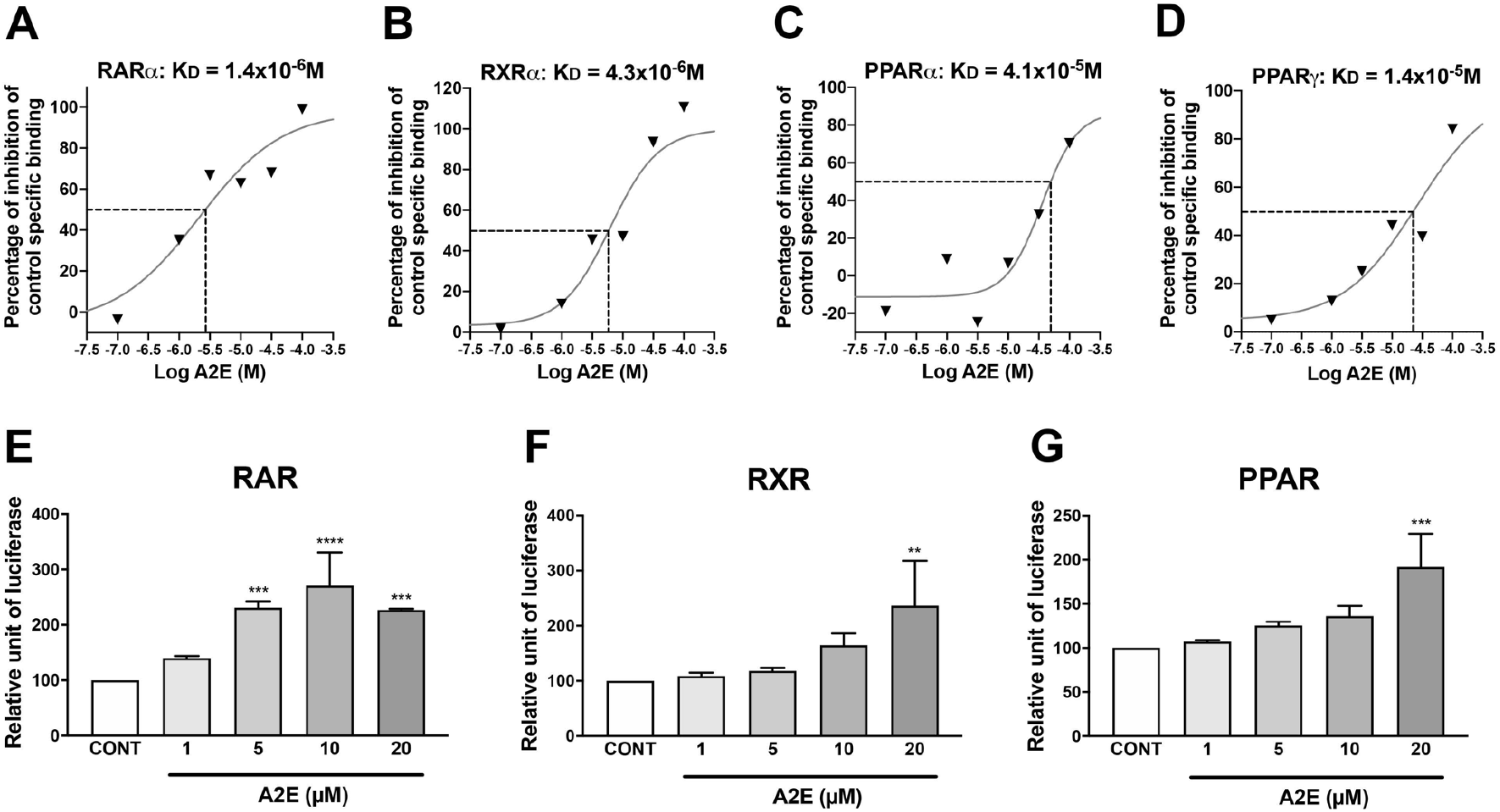
A2E binds RAR-α, RXR-α, PPAR-α and -γ, and transactivates RARs, RXRs, and PPARs. A2E binding to RAR-α (A), RXR-α (B), PPAR-α (C) and PPAR-γ (D) are represented. RARs (E), RXRs (F) and PPARs (G) transactivation induced by increasing A2E concentration. Bars represent mean ± s.e.m. with n=3. ** p < 0.01, *** p < 0.001, **** p < 0.0001 compared to control (CONT) (One-way ANOVA, Dunnett’s post-test).

**Table 1:**
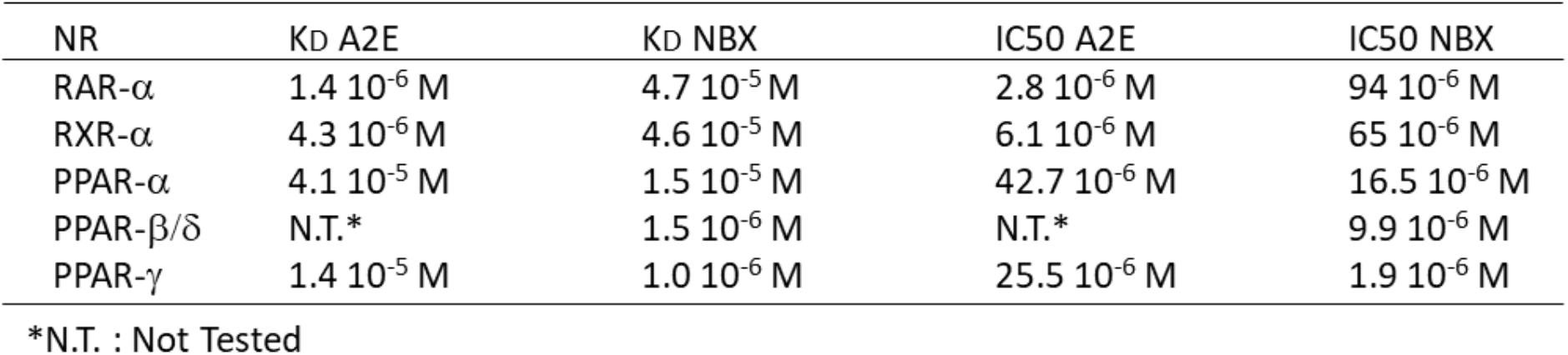
Respective Kd and IC50 for NRs of A2E and NBX

### Norbixin partially inhibits RXRs but enhances RARs transactivation induced by A2E

As A2E has been shown to be an agonist ligand of RAR-α (11), we hypothesized that norbixin might counteract A2E toxic effects by interacting with RAR-α and thus modulate RARs transactivation. To determine whether norbixin is also a ligand of RAR-α, binding affinity to RAR-α was investigated by competition experiments *in vitro* between norbixin and [^3^H] ATRA. We observed that norbixin binds to RAR-α but with a very weak affinity demonstrated by an IC50 of 94 10^−6^ M and a Kd of 4.7 10^−5^ M (Fig. 2A and Table 1). Next, we wanted to determine if norbixin modulates RARs transactivation induced by A2E. We compared the transactivation of RARs in primary porcine RPE cells in presence of A2E (20 μM) and in absence or presence of norbixin (20 μM). Norbixin did not inhibit, but on the contrary, it significantly increased A2E-induced RARs transactivation. Importantly, norbixin did not show any effect on RARs activation when added alone (Fig. 2B), and did not show any effect on RAR-α transactivation induced by BMS753 a specific high affinity agonist of RAR-α (Fig. 2C). Next, we wished to determine whether norbixin interferes with A2E-induced transactivation of RXRs. We first performed binding experiments by competition *in vitro* between norbixin and [^3^H] 9-*cis*-retinoic acid. We observed that norbixin binds RXR-α with a low affinity (IC50 of 65 10^−6^ M and Kd of 4.6 10^−5^ M) (Fig. 2D and Table 1). Then in transactivation experiments we observed that norbixin inhibited partially (by 49%, p < 0.01) the transactivation of RXRs induced by A2E, but had no effect when used alone (Fig. 2E). By contrast, norbixin did not inhibit the transactivation of RXRs induced by HX630 a selective high affinity pan RXRs agonist (Fig. 2F). In conclusion, our results show that norbixin binds RAR-α and RXR-α and potentiates RARs transactivation induced by A2E but has no effects when added with a high affinity RAR-α ligand. In addition, norbixin behaves as a partial neutral antagonist of RXRs transactivation induced by A2E, but has no effects on RXRs transactivation induced by a high affinity RXRs ligand.

**Figure 2.**
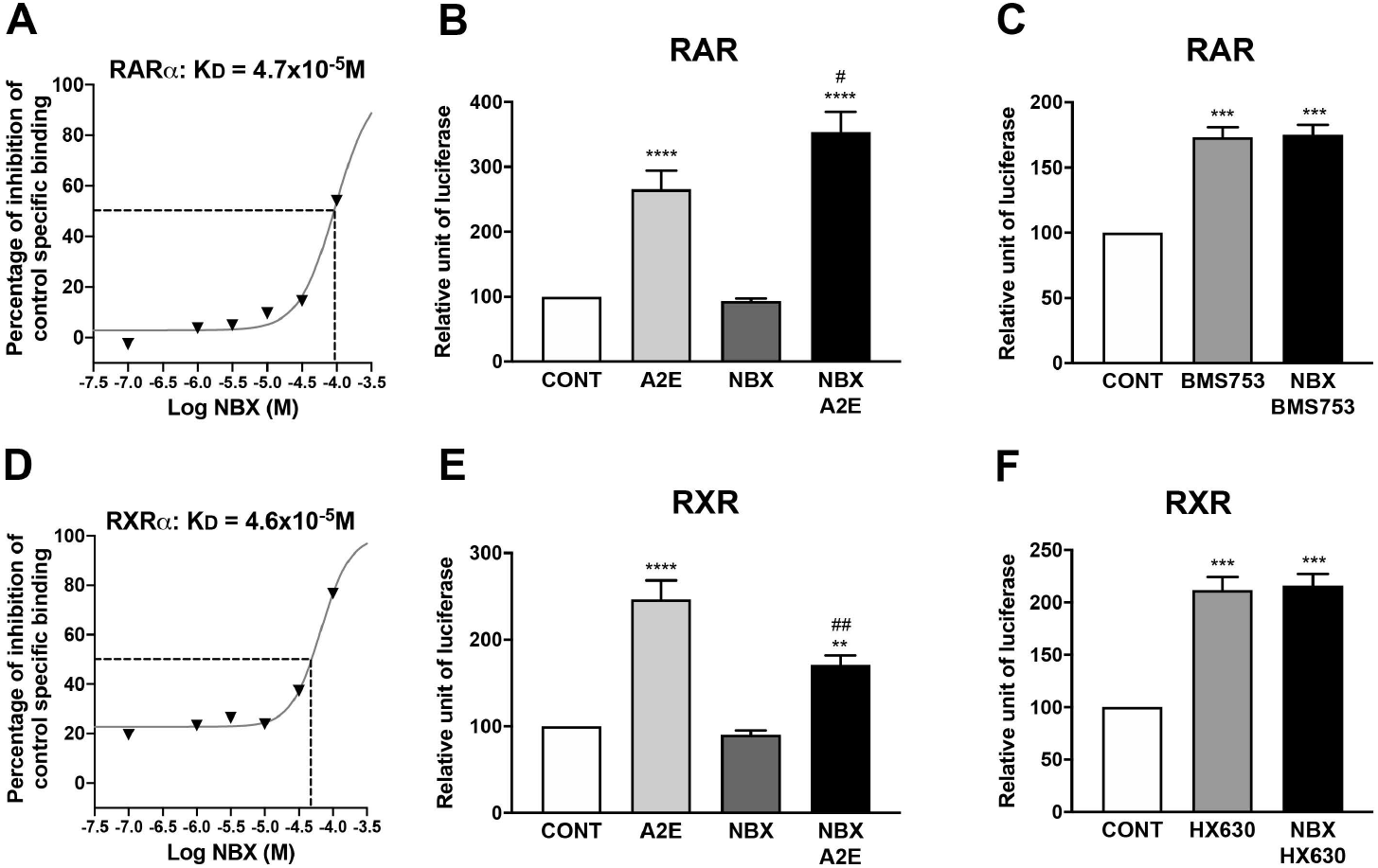
Norbixin partially inhibits RXRs but enhances RARs transactivation induced by A2E. Norbixin (NBX) binding to RAR-α (A) and RXR-α (D) are represented. Effect of A2E (20 μM), NBX (20 μM) alone and NBX (20 μM) + A2E (20 μM) on RAR (B) and RXR (E) transactivation. Effect of BMS753 (20 μM), a RAR (-α specific) agonist, and of HX630 (5 μM), a (pan) RXR agonist, alone or in competition with NBX (20 μM) on RAR (C) and RXR (F) transactivation. Bars represent mean ± s.e.m. with n=3-6. ^#^p < 0.05, ** or ^##^ p < 0.01, ***p < 0.001, *** p < 0.0001 compared to CONT or to A2E respectively (Oneway ANOVA, Dunnett’s post-test).

### Norbixin inhibits PPARs transactivation induced by A2E and inhibits PPAR-γ transactivation induced by troglitazone but not PPAR-α and -β/δ transactivation induced by high affinity agonists

We have shown here that A2E is a ligand of PPARs and induces their transactivation. It has been shown by *in silico* studies that norbixin is also a ligand of PPAR-γ (25). We wished to determine whether norbixin could modulate PPARs transactivation induced by A2E. Firstly, we performed competition experiments *in vitro* between norbixin and an agonist of PPAR-α, [^3^H] GW7647 and an agonist of PPAR-γ [^3^H] rosiglitazone. We report for the first time that norbixin binds to PPAR-α with an IC50 of 16.5 10^−6^ M (Fig. 3A) and a Kd of 1.5 10^−5^ M (Table 1). However, norbixin preferentially binds to PPAR-γ (IC50 =1.9 10^−6^ M and Kd = 1.0 10^−6^ M) (Fig. 3B and Table 1). Therefore, we tested whether norbixin would induce the transactivation of PPAR-α, -β/δ, or -γ by cotransfecting porcine RPE cells with a plasmid expressing each of the full length PPAR isoforms and a plasmid expressing luciferase under the control of the PPAR response element. While specific synthetic PPAR-α, -β/δ and -γ agonists (GW9578, GW0742 and troglitazone (TGZ) respectively) induced transactivation of the corresponding PPAR isoform (Fig. 3C, D, and E respectively), norbixin did not demonstrate any transactivation activity of its own on any PPARs (Fig. 3C to E). The same results were obtained following co-transfection with an expression plasmid for fusion protein of GAL4DNA-binding domain and PPAR ligand binding domain and with a plasmid expressing luciferase under the control of the GAL4 upstream activating sequences (data not shown). Interestingly, using an *in vitro* antagonist assay we also observed the antagonist activity of norbixin on PPAR-β/δ with an IC50 of 9.9 10^−6^ M and an affinity of 1.5 10^−6^ M (Table 1). Then, we wanted to determine if norbixin modulates the PPARs transactivation induced by A2E. We observed that norbixin inhibits the transactivation of endogenous PPARs induced by A2E in primary porcine RPE cells by 52%, (p < 0.0001) (Fig. 3F). To further define if norbixin inhibited all 3 PPARs isoforms or was specific of only some of them, specific constructs coding for PPAR-α, β/δ, or -γ were overexpressed separately and co-transfected using a plasmid containing the luciferase reporter gene under the control of the PPARs response element in porcine RPE cells. The transactivation of each PPARs isoform in response to A2E in presence or in absence of norbixin was determined by measuring luciferase activity. In this system, A2E induced the transactivation of all three PPARs isoforms (Fig. 3G to H). We observed that norbixin abolished completely the transactivation of PPAR-α, PPAR-β/δ, and PPAR-γ when these are overexpressed (Fig. 3G, H, and I respectively). In contrast with the inhibitory effect of norbixin on PPARs induced transactivation by A2E, norbixin did not inhibit significantly the transactivation of PPAR-α and -β/δ induced by their high affinity PPARs ligands GW9578, and GW0742 at 10 nM and 1 nM respectively (Fig. 3J, and K). Importantly, by contrast norbixin partially inhibited PPAR-γ transactivation induced in presence of 1 μM of TGZ (Fig. 3L) but this effect was lost when TGZ was used at a higher concentration (10 μM, data not shown). Altogether our results show that norbixin “behaves” as a neutral pan antagonist of PPARs when their transactivation is stimulated by A2E which has a low affinity for PPARs. Moreover, according with its respective affinities for the 3 PPARs isoforms, norbixin has no inhibitory effect on the transactivation of PPAR-α and -β/δ, but inhibits PPAR-γ transactivation when it is induced by their respective synthetic selective PPARs high affinity agonists.

**Figure 3.**
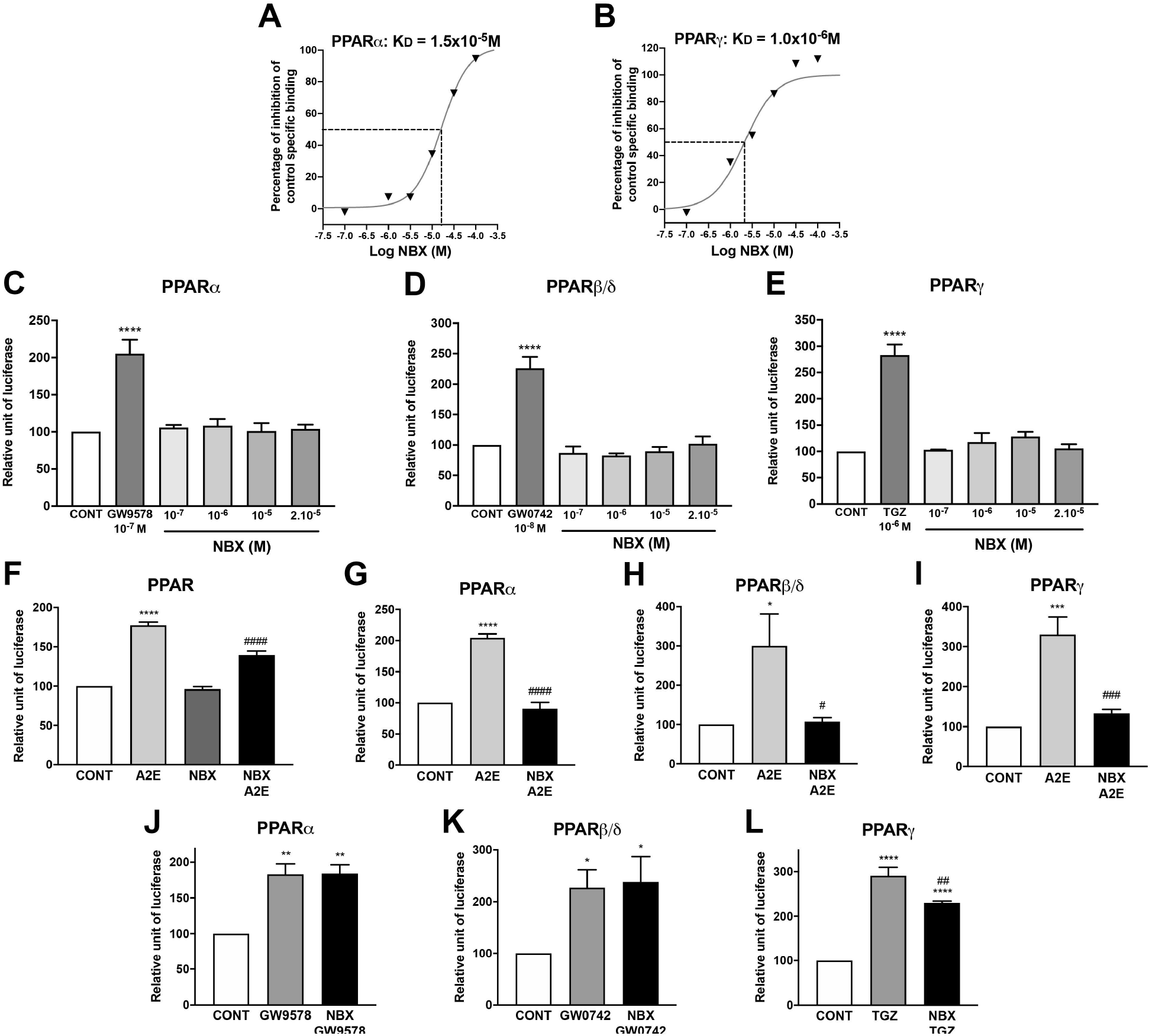
Norbixin inhibits PPARs transactivation induced by A2E and inhibits PPAR-γ transactivation induced by TGZ but not PPAR-γ and -β/δ transactivation induced by high affinity agonists. NBX binding to PPAR-α (A) and PPAR-γ (B) are represented. PPAR-α (C), PPAR-β/δ (D) and PPAR-γ (E) are not transactivated by NBX but by their own specific agonists (GW9578, GW0742 and TGZ) in RPE cells overexpressing each form of PPAR. Effect of A2E (20 μM), NBX (20 μM) alone and NBX (20 μM) + A2E (20 μM) on endogenous PPAR transactivation (F). Effect of A2E (20 μM) and NBX (20 μM) + A2E (20 μM) on over-expressed PPAR-α (G), PPAR-β/δ (H) and PPAR-γ (I) transactivation. Competition transactivation experiment between NBX and PPAR-α (J), PPAR-β/δ (K) and PPAR-γ (L) specific agonists (GW9578 (10 nM), GW0742 (1 nM), and TGZ (1 μM). Bars represent mean ± s.e.m. with n=3-6. * or ^#^p < 0.05, ** or ^##^ p < 0.01, *** or ^###^ p < 0.001, **** or ^####^ p < 0.0001; * compared to CONT; ^#^ compared to A2E (F-I) or to TGZ (L) (One-way ANOVA, Dunnett’s post-test).

### Norbixin regulates RPE survival and the expression of anti-apoptotic, inflammatory and angiogenic factors modulated by A2E

A2E toxicity on RPE cells and VEGF production has been specifically associated with RARs transactivation *in vitro* and *in vivo* (11, 12). Moreover, it has been shown that A2E might play a role in AMD pathogenesis by inducing inflammation (8–10) and the expression of VEGF *in vitro* and *in vivo*. RARs, RXRs and PPARs could be involved in these processes (14). We have shown that norbixin inhibits partially endogenous PPARs and RXRs transactivation and enhances the transactivation of RARs induced by A2E. Therefore, we aimed to decipher the effects of norbixin on A2E-induced expression of the anti-apoptotic related molecule Bcl2, inflammation and angiogenesis in RPE cells *in vitro*.

Firstly, we confirmed our previously published results (24) showing approximately 80% photoprotection of RPE cells by 20 μM of norbixin following blue light exposure in presence of A2E (Fig. 4A). It has been shown previously that Bcl2 protects against A2E and blue light induced apoptosis (27). We measured by western blot the expression of this protein in primary porcine RPE cells cultivated in presence and absence of A2E and norbixin. We showed that norbixin is able to maintain partially the level of Bcl2 expression decreased by A2E in RPE cells (Fig. 4B).

**Figure 4.**
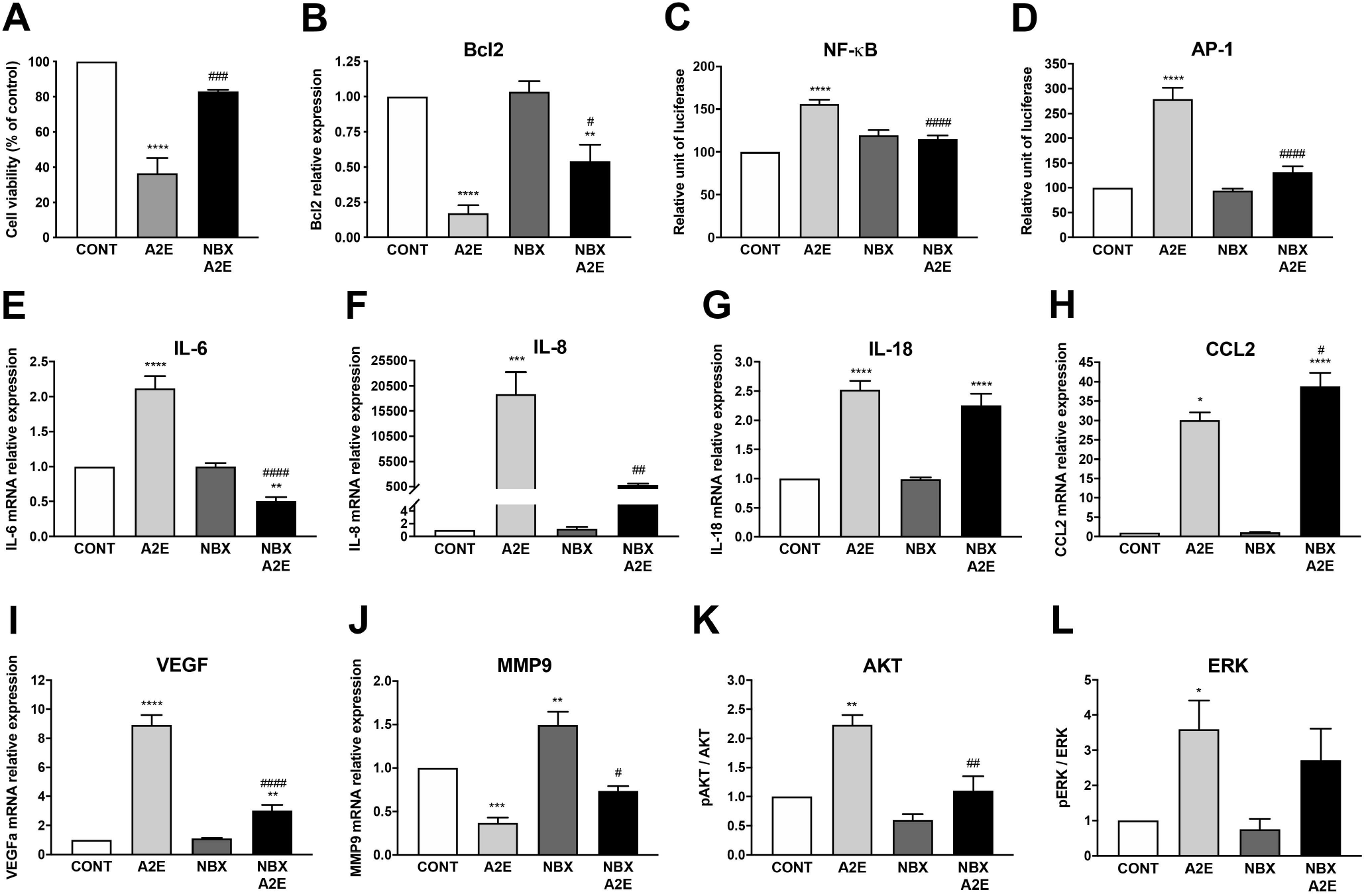
Norbixin regulates RPE survival and the expression of anti-apoptotic, inflammatory and angiogenic factors modulated by A2E. Effect of NBX (20 μM) on A2E-induced RPE cell culture phototoxicity model (A). Effect of A2E (30 μM), NBX (20 μM) alone and NBX (20 μM) + A2E (30 μM) on Bcl2 protein expression (B), on IL-6 (E), IL-8 (F), IL-8 (G), CCL2 (H), VEGF (I) and MMP9 (J) mRNA expression, and on AKT (K) and ERK (L) protein phosphorylation. Effect of A2E (20 μM), NBX (20 μM) alone and NBX (20 μM) + A2E (20 μM) on NF-κB (C) and AP-1 (D) transactivation. Bars represent mean ± s.e.m. with n=3-6. * or ^#^p < 0.05, ** or ^##^ p < 0.01, ***p < 0.001, **** or ^####^ p < 0.0001 compared to CONT or to A2E respectively (One-way ANOVA, Dunnett’s post-test).

Secondly, we compared the transactivation of transcription factors involved in inflammation regulation and the expression of inflammatory cytokines induced by A2E in presence or absence of norbixin. We showed that A2E induced the transactivation of NF-κB, a pivotal mediator of inflammatory responses regulating multiple aspects of innate and adaptative immune functions. NF-κB transactivation induced by A2E was strongly downregulated by norbixin (73%, Fig. 4C). A2E also induced transactivation of AP-1, a key modulator in inflammatory cytokine expression, which was also inhibited by norbixin (82%, Fig. 4D). Accordingly, norbixin was able to significantly reduce (145%) the mRNA expression of IL-6 induced by A2E (Fig. 4E). The level of IL-6 following treatment with norbixin was even lower than the basal level found in RPE cells non treated with A2E. We observed that norbixin strongly inhibited (approximately by 23-fold) IL-8 mRNA expression as well (Fig. 4F). It has been previously shown that A2E activates the NLRP3 inflammasome complex and induces the pro inflammatory IL-1β cytokine expression in RPE cells (8). In our *in vitro* model however, we were not able to detect significant levels of IL-1β mRNA and protein (data not shown). Nevertheless, we observed that mRNA expression of IL-18, a cytokine of the IL-1 family, also depending on NLRP3 activation and that has been implicated in AMD pathogenesis (28), was induced by A2E in RPE cells (Fig. 4G). In contrast with IL-6 and IL-8, IL-18 expression was not inhibited in the presence of norbixin (Fig. 4G). The expression of the chemokine CCL2, that is also implicated during AMD through macrophages recruitment in the subretinal space (29), was up-regulated by A2E (Fig. 4H). Unexpectedly its expression was not inhibited but was further increased by norbixin treatment (Fig. 4H). Altogether, norbixin displays differential modulatory effects on expression of antiinflammatory and pro-inflammatory molecules that might be related to its inhibitory effect on PPARs and RXRs.

Thirdly, angiogenesis is the major driver of neovascular AMD, during which VEGF promotes the growth of neo-vessels originating from the choriocapillaris towards the retina. In our *in vitro* model of porcine RPE cells we confirmed that A2E stimulation enhanced significantly (by approximately 9-fold) the mRNA expression of VEGF (Fig. 4I). However, we also observed that this increase in VEGF expression was significantly inhibited by addition of norbixin (Fig. 4I). VEGF alone is not sufficient to induce neovascular AMD if Bruch’s membrane remains intact (30). Matrix metalloproteinases and in particular MMP9 are able to alter Bruch’s membrane integrity (30), thus potentially allowing neo-vessels responsible for blindness in neovascular AMD to reach the neural retina responsible for blindness in neovascular AMD. Since MMP9 production has been shown to be under the control of PPAR-α (31), we measured MMP9 mRNA expression in our *in vitro* model of porcine RPE cells stimulated by A2E in presence or in absence of norbixin. Surprisingly, A2E reduced the expression of MMP9 mRNA (Fig. 4J), whereas norbixin slightly increased MMP9 expression in the absence or presence of A2E (Fig. 4J).

Finally, it has been shown that RPE cell damage following UVB exposition is mediated by AKT and ERK activation (32). Indeed, both AKT (33) and ERK (34) control apoptosis induction. In addition, it has been shown that ERK inhibition suppress VEGF production in RPE cells (34) and IL-6 production in fibroblasts (35). Thus, we reasoned that, since A2E induces apoptosis of RPE cells and IL-6 and VEGF production, ERK and AKT could be some targets of A2E and that their action could be regulated by norbixin. We tested the effects of norbixin on these signalling pathways in RPE cells treated with A2E. We observed that A2E indeed induced the phosphorylation of AKT (Fig. 4K) and of ERK (Fig. 4L) in primary porcine RPE cells. Treatment with norbixin abrogated completely the increase of phosphorylation of AKT (Fig. 4K). By contrast, despite a small trend towards reduction, ERK phosphorylation was not significantly inhibited by norbixin (Fig. 4L).

## Discussion

It has been previously shown that A2E is a ligand of RAR-α and it was proposed that RPE cell toxicity *in vivo* and VEGF expression *in vitro* and *in vivo* induced by A2E are mediated through RARs transactivation (11,12). Here we define the profile of A2E binding affinities for RAR-α, RXR-α and all three PPARs isoforms. Importantly, in the present study we confirm that A2E is a retinoid binding to RAR-α and inducing the transactivation of RARs. But we also show for the first time that A2E binds to and induces the transactivation of RXRs and of all three isoforms of PPARs.

Based on *in silico* and *in vivo* studies it had been previously reported that norbixin is a PPAR-γ agonist such as pioglitazone (25). However, NRs responsiveness in response to ligands depends on the specific cellular context and experimental settings (16, 19). Indeed, by contrast, it has been shown that norbixin at 20 μM did not induce PPAR-γ transactivation in 3T3-L1 adipocytes (26). In the present study, we specified, in the context of primary porcine RPE cells, the interactions between norbixin and PPARs, RAR-α and RXR-α and the effects of norbixin on PPARs, RARs and RXRs transactivation. By competitive binding experiments using radiolabelled ligands, we showed that norbixin is not only a ligand of PPAR-γ, but also of PPAR-β/δ (data not shown) and of PPAR-α. Moreover, norbixin also binds, albeit with low affinity, to RXR-α and RAR-α. However, despite positive binding experiments, norbixin alone in our experimental settings was unable to induce the transactivation of any of the NRs tested. By contrast, norbixin inhibited the transactivation induced by A2E of RXRs and all three isoforms of PPARs. In addition, norbixin enhanced RARs transactivation induced by A2E. These observations suggest that norbixin is a neutral antagonist of RXRs and PPARs. However, importantly we also showed that norbixin did not inhibit the transactivation of PPAR-α and -β/δ isoforms stimulated by high affinity synthetic agonists (GW9578, and GW0742 respectively), and norbixin only partially inhibited PPAR-γ transactivation induced by low (1 μM) but not higher (10 μM) concentrations of TGZ. Thus, the inhibitory effect of norbixin on PPARs transactivation is observed when it is induced by A2E, which has a low affinity for PPARs, but not when it is induced by high affinity synthetic PPARs ligands. The respective affinities of the various PPARs ligands compared to the affinity of norbixin considered as a competitor might explain this apparently contradictory results. Indeed, we showed that A2E binds PPAR-γ and PPAR-α with low affinities (Kd of 1.4 10^−5^ M and 4.1 10^−5^ M, respectively) that are inferior to the affinities of norbixin for PPAR-γ (Kd of 1.0 10^−6^ M) and PPAR-α (Kd of 1.5 10^−5^ M). Therefore, norbixin appears able to displace A2E from the three PPARs isoforms. By contrast norbixin might not be able to compete with the synthetic high affinity PPARs agonists GW9578 and GW0742, whose binding affinities are in the nanomolar range, and only weakly with TGZ, whose binding affinity has been reported to be 1 μM [11]. Thus, norbixin “behaves” as a neutral pan PPARs antagonist when their transactivation is induced by A2E. However, norbixin might not be considered strictly speaking as a PPARs antagonist because it doesn’t inhibit PPARs transactivation induced by high affinity agonists. Norbixin should rather be considered as an inhibitor of PPARs transactivation, when it is induced by low affinity ligands such as A2E.

Due to the permissive nature of PPARs heterodimers (16, 19), A2E-induced transactivation of PPAR/RXR heterodimers could result from direct binding of A2E to PPARs, RXR-α or both. Moreover, it has been shown that an agonist ligand of RXRs/RXRs homodimers can also induce the transactivation of PPARs by direct binding on the PPARs response elements (36). Therefore, it could be hypothesized that inhibition of A2E-induced PPARs transactivation by norbixin may result not only from the direct binding of norbixin on PPARs, as demonstrated above, but also by an indirect effect through binding of norbixin to RXR-α. Accordingly, here we observed the partial inhibition of A2E-induced transactivation of RXRs by norbixin. This inhibitory effect of norbixin could also be explained by their respective binding affinities to RXR-α. Indeed, the binding affinity of A2E for RXR-α is 10-fold higher than the binding affinity of norbixin (Kd of 4.3 10^−6^ M versus 4.6 10^−5^ M respectively). Therefore, norbixin could only partially compete with A2E for RXR-α. Similar to our observations that norbixin is unable to, or only partially, compete with selective high affinity PPARs agonists, we show that norbixin is unable to inhibit the transactivation of RXR-α induced by HX630 a high affinity pan agonist of RXRs. The interactions between A2E and norbixin with RXR-β and -γ were not evaluated here but should also be taken into account to explain the incomplete inhibition by norbixin of A2E-induced RXRs transactivation. Finally, we showed that A2E binds to RAR-α with a Kd of 1.4 10^−6^ M while affinity of norbixin for RAR-α is much lower (Kd of 4.7 10^−5^ M). It is therefore not surprising that norbixin is unable to inhibit RARs transactivation induced by A2E nor RARs transactivation induced by BMS753, a high affinity selective RAR-α agonist. By contrast, we observe that norbixin enhances the transactivation of RARs induced by A2E. Since norbixin alone has no effect on RARs transactivation and that norbixin may displace A2E from PPARs and RXRs, we make the hypothesis that the increased RARs transactivation by norbixin in the presence of A2E is an indirect effect. Indeed, free A2E that could no longer bind to PPARs and RXRs would be able to further transactivate RARs. Nevertheless, here again the importance of the RAR-β and -γ isoforms in norbixin effects on A2E induced RARs transactivation deserves further investigations.

PPARs ligands, retinoids and rexinoids, through their binding to NRs, exert wide effects on cell survival, cytokines production regulation, and angiogenesis (14) which are all associated with AMD pathophysiology (20–22). The modulation by norbixin of RARs, RXRs and PPARs transactivation induced by A2E suggests a role of these NRs in mediating A2E deleterious biological effects. For instance, our results suggest that norbixin-induced photoprotection of primary porcine RPE cells from A2E and blue light illumination (24) through maintenance of Bcl2 expression, (27) is regulated via RARs, RXRs and/or PPARs transactivation. Similarly, these NRs appear crucial in regulation of inflammation induced by A2E as demonstrated by the modulatory effect of norbixin on A2E-induced expression of IL-6, IL-8, MMP9, and CCL2 as well as on the transactivation of NF-κB and of AP-1, and on the phosphorylation of AKT. In addition, our observations suggest that A2E-induced expression of the angiogenic factor VEGF could also be under the control of RARs, RXRs and/or PPARs transactivation. By contrast, regulation of IL-18 expression and ERK-phosphorylation appear independent of RARs, RXRs and PPARs regulation. However, this doesn’t preclude that IL-18 expression and ERK-phosphorylation might be modulated via other NRs.

The fact that A2E induces RXRs transactivation implies that A2E may also induce the transactivation of other permissive NRs such as LXRs that have recently been reported to play an important role in AMD (39). Indeed, LXRs are involved in regulation of inflammation, angiogenesis and cell survival. It has been demonstrated that LXRs regulate, at least partially, the expression of VEGF (37) and MMP9 (38), and here we demonstrated that their expressions are modulated by A2E and also by norbixin. Moreover, LXRs have been implicated in the regulation of APOE and ABCA1 (39) that are important for the efflux of cholesterols internalized during daily uptake of photoreceptor tip cells. Since A2-PE and other A2E precursors, which are present in photoreceptor outer segments, are internalized during the same process, it could be hypothesized that LXRs may also play a role in the metabolism of A2E by RPE cells. We have shown previously that norbixin reduces A2E accumulation *in vitro* (24). The reduction of intracellular concentrations of A2E may affect the level of transactivation of RARs, RXRs and PPARs. The analysis of the relations existing between A2E accumulation, NRs transactivation and their modulation by norbixin *in vitro* and *in vivo* may help better understand AMD pathophysiology. Thus, the precise effects of A2E and norbixin on LXRs will be the subject of future experiments.

Nevertheless, the regulation by NRs of molecules involved in inflammation and angiogenesis remains complex. For instance, it has been shown that ATRA, a RARs agonist, suppress IL-6 independently of RARs activation but through ERK inhibition (35). By contrast, in our experiments, norbixin suppression of IL-6 expression induced by A2E was associated with enhanced RARs transactivation but not with modulation of ERK phosphorylation. Furthermore, we showed that norbixin inhibits transactivation of NF-κB and AP-1 transcription factors, that are essential for induction of inflammation and of IL-6 in particular. Redundancy between several signalling pathways complexifies the understanding of cytokines regulation. The same biological effect might be observed but the molecular events leading to it might differ and upon cell environment, one signalling pathway may be prepondering upon another. A second example of this complexity is given by MMP9 regulation. It has been reported that, in addition to LXRs, PPAR-α or -γ activation might be responsible for MMP9 downregulation in macrophages following LPS stimulation (40). Our experiments support this assumption as we show that A2E induced the transactivation of both these PPARs isoforms and reduced MMP9 expression. Accordingly, treatment with norbixin restored but only partially MMP9 levels in presence of A2E potentially through inhibition of PPAR-α and -γ transactivation. However, concomitantly norbixin inhibited A2E-induced transactivation of NF-κB and AP-1 transcription factors, which are required for MMP9 induction following LPS treatment (31, 40, 41). Thus, inhibition of A2E-induced transactivation of NF-κB, AP-1 and PPARs by norbixin has potentially opposing effects on MMP9 expression. Alternatively, it has also been reported that ATRA inhibits MMP9 expression and activity in a human breast cancer cell line (42). Therefore, RARs induced transactivation by A2E could explain the inhibition of MMP9 expression by A2E and why norbixin that enhances RARs transactivation by A2E cannot restore completely MMP9 expression. However, none of these signalling pathways could explain the increase of MMP9 expression observed in RPE cells treated with norbixin alone. This puzzling observation suggests that other transcriptions factors, or NRs such as LXRs might be involved as well in MMP9 expression regulation. Regarding CCL2 expression, its regulation by A2E and norbixin appears different from other molecules studied in the present article. Indeed, while CCL2 expression was induced by A2E, it was further enhanced by norbixin treatment. It has been shown that 9-*cis*-RA, a bi-specific agonist of RARs and of RXRs induces CCL2 expression in THP-1 cells, a human monocytic cell line (43). Interestingly, it has also been reported that an agonist of PPAR-α inhibits CCL2 production in an *in vivo* model of corneal alkali burn (44). Norbixin by suppressing A2E-induced PPAR-α transactivation may thus release A2E inhibitory effect through PPAR-α resulting in increased CCL2 production. However, antagonizing PPAR-γ also reduces CCL-2 expression during microglia differentiation *in vitro* (45). Altogether, these conflicting observations suggest that, upon A2E treatment, CCL2 overexpression may result from a subtle equilibrium between RARs, PPAR-α and -γ transactivation levels. Alternatively, as proposed above, it could be hypothesized that a competition for A2E between the various NRs (RARs, RXRs and PPARs) exists. In this hypothesis, norbixin, by impairing the binding of A2E to PPARs and RXRs, could enhance A2E binding to RARs and their transactivation, thus promoting CCL2 expression. This may indeed be the case, because we showed that norbixin increased A2E-induced RARs transactivation. Further experiments are required to decipher the exact mechanism of CCL2 upregulation by A2E and norbixin. The potential role of RARs, RXRs and PPARs in the regulation of inflammatory cytokines, pro-angiogenic, and antiapoptotic molecules by A2E and norbixin is schematically summarized in figure 5.

**Figure 5.**
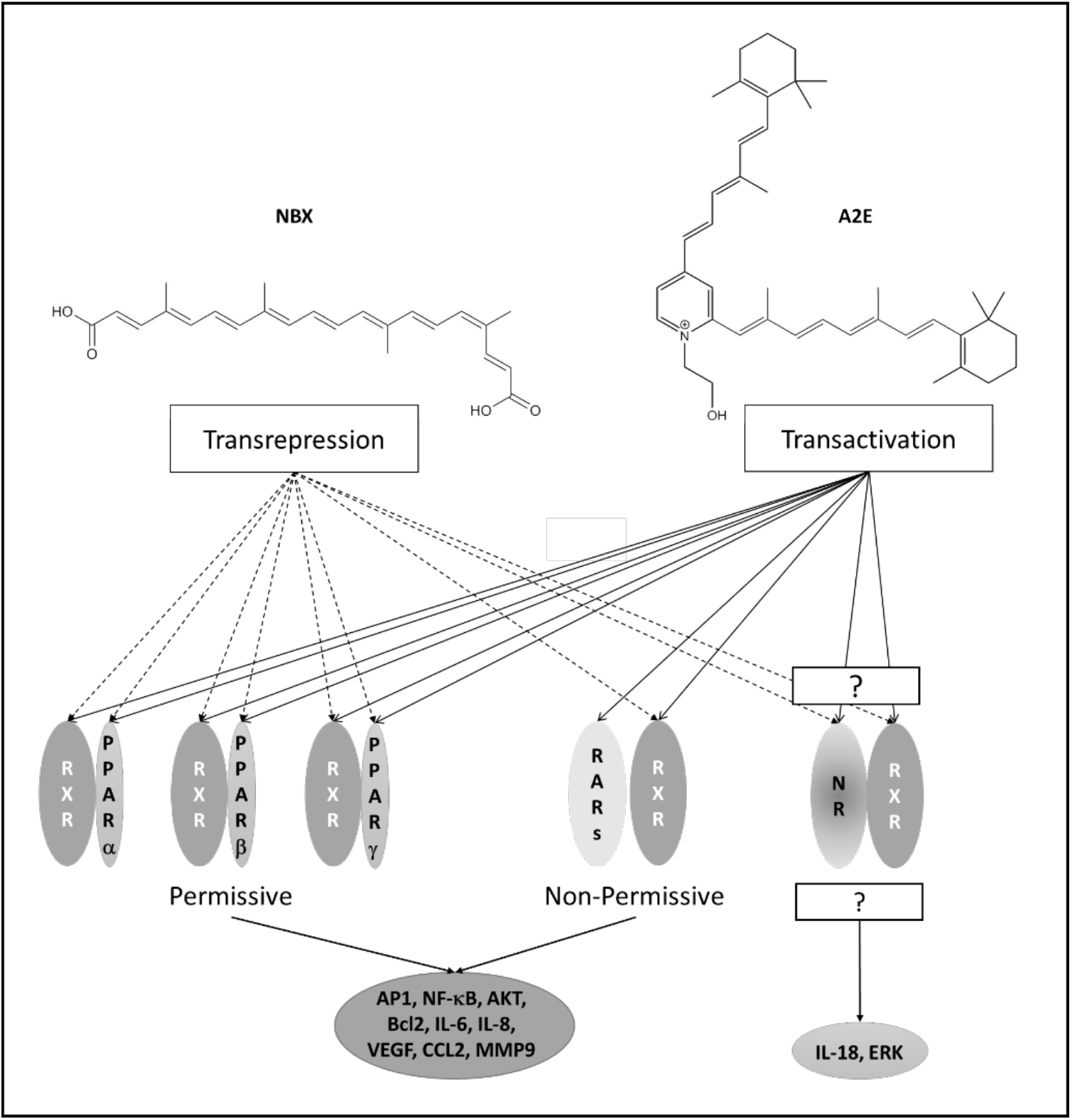
Summary of interactions of norbixin and A2E on NRs and their roles in genes expression regulation. A2E is a ligand of RAR-α, RXR-α and of PPARs and induces their transactivation while NBX trans-represses PPARs and RXRs but enhances RARs transactivation in presence of A2E. The permissive/non permissive status of the nuclear receptors (NRs) is represented. NBX inhibits AP-1, and NF-κB transactivation, inhibits the expression of the inflammatory cytokines (IL-6 and IL-8) and of the angiogenic molecule VEGF, and reduces the phosphorylation of AKT while enhancing protein level of the anti-apoptotic molecule Bcl2 and CCL2 and MMP9 mRNA expression. By contrast NBX has no effect on IL-18 expression and ERK phosphorylation suggesting (an)other(s) NR(s) might be involved in their regulation.

Altogether, here we report that A2E effects on RPE cell death, cytokine regulation, and angiogenesis, which are all important during AMD pathophysiology, may be controlled at least partly through RARs, RXRs and PPARs modulation. We are not the firsts to point out the potential importance of NRs and particularly, PPARs in controlling several AMD-pathogenic pathways (46–51). Indeed, it has been shown that PPAR-α agonists protect RPE cells from an oxidative stress insult *in vitro* (52) and reduce angiogenesis in mouse models of neovascular AMD (53). PPAR-β/δ modulation seems to play different roles in dry AMD and neovascular AMD models. Pharmacological inhibition of PPAR-β/δ signalling *in vitro* on RPE cells or *in vivo* in the mouse laser-induced choroidal neovascularization model, limited angiogenesis (54), but exacerbated signs of dry AMD. Conversely, activation of PPAR-β/δ reduced lipid accumulation in RPE cells *in vitro* (a symptom in intermediate dry AMD) [55]. In parallel, it has been shown that agonists of PPAR-γ such as TGZ and 15-deoxy-Δ^12,14^-prostaglandin J2 protect RPE cells from an oxidative stress insult (55, 56), whereas other PPAR-γ agonists (pioglitazone and rosiglitazone) exacerbate cell death induced by t-butyl hydroperoxide treatment (56). Moreover, PPAR-γ activation by TGZ reduces VEGF production in RPE cells, reduces tube formation in choriocapillaris endothelial cells *in vitro* and reduces angiogenesis in a rat model of laser-induced choroidal neovascularization (57). Despite the fact that they may appear to contradict some of the studies cited above, our results are novel and important because they demonstrate for the first time that A2E, which plays a central role in AMD, behaves itself as a PPARs pan agonist as well as an RXRs and RARs agonist. Moreover, we report that norbixin may regulate A2E activity by antagonizing partially PPARs and RXRs and enhancing indirectly RARs transactivation. These observations of potentially beneficial effects of norbixin *in vitro* are consistent with our previous published results showing that norbixin is indeed protective in various *in vivo* animal models of AMD ((24) and Fontaine et al. 2020, article in press).

In conclusion, we show for the first time that A2E induces the transactivation of RXRs and PPARs in addition to RARs transactivation, and that an important part of the biological effects of A2E may be mediated through these NRs activation. Moreover, we demonstrate that norbixin partially inhibits RXRs transactivation and behaves as a pan inhibitor of PPARs transactivation induced by A2E. Moreover, norbixin enhances A2E-induced RARs transactivation potentially via an indirect effect. Finally, here we show that norbixin modulates the expression of molecules involved in angiogenesis, inflammation and apoptosis, induced by A2E and that are critical for AMD pathogenesis. Our study opens new avenues for the treatment of AMD using norbixin or related molecules.

## Experimental procedures

### Reagents/Chemicals

All usual chemicals and primers were from Sigma (St. Louis, MO, USA). Reagents for cell culture, transfection, and quantitative RT-PCR were from Thermo Fisher Scientific (Waltham, MA, USA). RNA extraction NucleoSpin^®^ RNA kit was from Macherey Nagel (Düren, Germany). ECL prime and PVDF membrane were from Amersham GE Healthcare (Buckinghamshire, UK). Primary antibodies against the following proteins were used: GAPDH (Santa Cruz Biotechnology, Inc, Dallas, TX, USA); Bcl2 (BD Biosciences, Franklin Lakes, NJ, USA); pAKT, AKT, pERK and ERK (Cell Signaling, Danvers, MA, USA). Secondary antibodies were from Jackson ImmunoResearch (Cambridgeshire, UK). BMS753 was from Santa Cruz, GW9578 was from Cayman Chemical Company (Ann Arbor, MI, USA) and GW0742, TGZ and HX630 were purchased from TOCRIS (Bristol, UK). Cignal Pathway Reporter Assay Kits were from QIAGEN (Frederick, MD, USA). The Dual-Luciferase Reporter Assay System was purchased from Promega (Madison, WI, USA). pcDNA 3.1 (+)-PPARα and pcDNA 3.1 (+)-PPARβ/δ (pig sequence) were purchased from Genscript (Piscataway, NJ, USA). pCMV6-XL4-PPARγ (human sequence) was purchased from Origene (Rockville, MD, USA).

### Synthesis of norbixin

9’-*cis*-norbixin was prepared from 9’-*cis*-bixin (AICABIX P, purity 92%) purchased from Aica-Color (Cusco, Peru) upon alkaline hydrolysis as previously described (24) and according to Santos et al. (58). The obtained product (the 9’-*cis* isomer) showed an HPLC purity of 97% as confirmed by ^1^H-nuclear magnetic resonance (using malonic acid as internal standard). Fresh solutions of 9’-*cis*-norbixin stored in powder at −80°C were prepared in DMSO.

### Synthesis of A2E and A2E-Propylamine

A2E was synthesized by Orga-link (Magny-Les-Hameaux, France) as described before (59). Briefly, all-*trans*-retinal, ethanolamine and acetic acid were mixed in absolute ethanol in darkness at room temperature over 7 days. The crude product was purified by preparative HPLC in the dark to isolate A2E with a purity of 98% as determined by HPLC. A2E (20 mM in DMSO under argon) was stored at −20°C.

### In vitro model of RPE phototoxicity and treatments

Pig eyes were obtained from a local slaughterhouse and transported to the laboratory in icecold Ringer solution. After removal of the anterior segment of the eye, the vitreous and neural retina were separated from the RPE and removed. The eyecup was washed twice with PBS, filled with trypsin (0.25% in PBS) and incubated at 37°C for 1.5 h. RPE cells were harvested by gently pipetting, centrifuged to remove trypsin and re-suspended in Dulbecco’s Modified Eagle Medium (DMEM) supplemented with 20% (v/v) foetal-calf serum (DMEM 20%FCS) and 0.1% gentamycin. Cells were seeded into 60 mm diameter Petri dishes, cultured in an atmosphere of 5% CO2/95% air at 37°C, and supplied with fresh medium after 24 h and 4 days *in vitro*. After one week in culture, cells were trypsinized and transferred to a 96-well plate at a density of 1.5 x 10^5^ cells/cm^2^ in DMEM 2%FCS. After 2 days *in vitro*, A2E was added to the medium at a final concentration of 30 μM, and 19 h later blue-light illumination was performed for 50 min using a 96 blue-led device (Durand, St Clair de la Tour, France) emitting at 470 nm (1440 mcd, 8.6 mA). Just before illumination, the culture medium was replaced by a modified DMEM without any photosensitizer and with 2% FCS. 24 h after blue-light irradiation, all cell nuclei were stained with Hoechst 33342 and nuclei of dead cells with ethidium homodimer 2, fixed with paraformaldehyde (4% in PBS, 10 min) and 9 pictures per well were captured using a fluorescence microscope (Nikon TiE) equipped with a CoolSNAP HQ2 camera and driven by Metamorph Premier On-Line program. Quantification of live cells was performed using Metamorph Premier Off-Line and a home-made program by subtraction of dead cells from all cells. Cell treatments were performed as followed. All the drugs used in these experiments were prepared as stock solutions in DMSO. Norbixin was added to the culture medium 48 h before illumination. For experiments aimed at measuring mRNA or protein expression in RPE, cells were seeded in 24-well or 6-well plates and the treatments were performed as before but the experiment was stopped before illumination. For mRNA analysis cells were lysed using the lysis buffer from the NucleoSpin^®^ RNA kit, and the sample was stored at −80°C. For protein analysis cells were collected in Eppendorf tubes, frozen in liquid nitrogen and kept at −80°C.

### Binding studies to RAR-α, RXR-α and PPAR-α and γ

Binding studies of A2E and norbixin to RAR-α, RXR-α and PPAR-α and γ were performed *in vitro* by an external laboratory (Eurofins Cerep, Celle L’Evescault, France) through competition experiments between A2E or norbixin and [^3^H] ATRA the natural ligand of RAR-α, [^3^H] 9-*cis*-retinoic acid as the natural ligand of RXR-α, [^3^H] GW7647 an agonist of PPAR-α and [^3^H] rosiglitazone an agonist of PPAR-γ.

### RAR, PPAR, RXR, AP-1 and NF-κB transactivation assays

After one week in culture, cells were trypsinized and transferred to a 96-well plate at a density of 6 x 10^4^ cells/cm^2^ in DMEM2%FCS. The next day, cells were transfected using the Cignal Reporter Assay Kits for RARs, PPARs, RXRs, AP-1 and NF-**κ**B, according to the manufacturer’s specifications. To measure the specific activation of PPARs isoforms, cells were also co-transfected with PPAR-α, PPAR-β/δ or PPAR-γ expression vectors. Transfection was performed with Lipofectamine and Plus Reagent in serum-free medium. 3 h after the transfection, the medium was replaced and treatments with the molecules to assay were performed. Luciferase activity was measured the next day using the Dual-Luciferase Reporter Assay System. The measurements were performed with a luminometer (Infinite M1000 from Tecan, Mannedorf, Switzerland). Firefly:Renilla activity ratios were calculated for each condition and ratios from transcription factor-responsive reporter transfections were divided by ratios from negative control transfections to obtain relative luciferase unit, as described by the manufacturer. At least 3 independent transfections were performed in triplicate for each condition.

### Quantitative RT-PCR

Total RNA was extracted using the NucleoSpin^®^ RNA kit according to manufacturer’s instructions. RT of 500 ng of RNA was performed using the SuperScript III Reverse Transcriptase following the manufacturer’s instructions. Five ng of cDNA were amplified using the SYBR GREEN realtime PCR method. PCR primers for target genes and housekeeping gene GAPDH were designed using Primer3Plus Bioinformatic software (Table 2). The RT-PCR using the StepOne Plus (Life Technologies, Carlsbad, CA, USA) consisted of incubation at 50°C for 5 min followed by 40 cycles of 95°C for 15 s and of 60°C for 1 min. The reaction was completed by a melt curve stage at 95°C for 15 s, 60°C for 1min, 95°C for 15 s and a final step at 60°C for 15 s. The relative mRNA expression was calculated using the comparative threshold method (Ct-method) with GAPDH for normalization. All experimental conditions were processed in triplicate and each experiment was done at least 3 times.

**Table 2:**
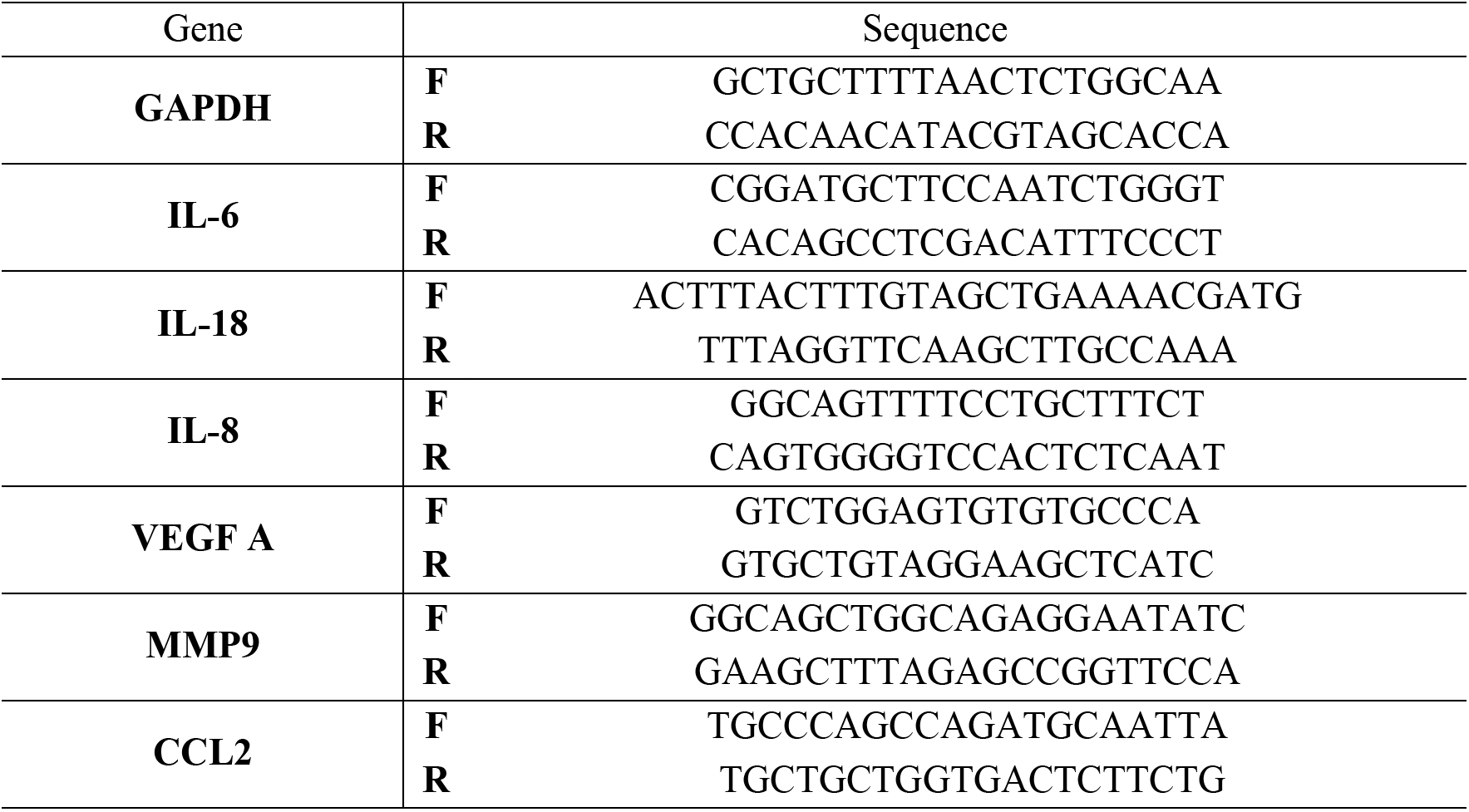
probes used for mRNA quantification by RT-QPCR

### Protein analysis

RPE samples were lysed in 20 mM Tris-HCl, 150 mM NaCl, 1 mM EDTA, 1% NP-40, pH 7.5 buffer containing a cocktail of protease and phosphatase inhibitors. The equal amount of proteins was resolved by 12% SDS polyacrylamide gel electrophoresis and electrotransferred onto a PVDF membrane using a standard protocol. The membranes were blocked with 5% milk or 10% BSA for 1 h, followed by incubation with a primary antibody overnight at 4°C. Subsequently the membranes were incubated with the corresponding horseradish peroxidase-conjugated secondary antibodies for 1 h. The signal was developed using enhanced chemiluminescence reagents ECL prime detection kit, quantified by densitometry using Bio1D (Vilber Lourmat, Germany) and normalized by GAPDH levels. Each experiment was done at least 3 times.

### Statistical analyses

For statistical analyses one-way ANOVA followed by Dunnett’s tests were performed using Prism 7 (GraphPad Software, La Jolla, CA, USA).

## Data availability

A patent “Composition for the protection of retinal pigment epithelium” covering the topic of this manuscript has been filed on April 30, 2015 (FR 15 53957) and is owned by Stanislas Veillet, René Lafont, José-Alain Sahel, Valérie Fontaine. Stanislas Veillet is founder, CEO and shareholder of Biophytis. René Lafont is founder, and shareholder of Biophytis. This does not alter our adherence to The Journal of Biological Chemistry policies on sharing data and materials. We have no restriction to share these data.

## Acknowledgements

The contribution of Dr L.N. Dinan for critical reading of the manuscript and language improvement is acknowledged.

## Abbreviations

(A2E): N-retinylidene-N-retinylethanolamine
(A2-PE): phosphatidylethanolamine-bisretinoid
(AKT): protein kinase B
(AMD): age-related macular degeneration
(AP-1): activator protein 1
(ATRA): all-*trans*-retinoic acid
(Bcl2): B-cell lymphoma 2
(CCL2): C-C motif chemokine ligand 2
(ERK): extracellular signal-regulated signal
IL: (interleukin)
(LXRs): liver X receptor
(NBX): norbixin
(NRs): nuclear receptors
(MMP9): matrix metalloproteinase 9
(PPARs): peroxisome proliferator activated receptors
(RARs): retinoic acid receptors
(RPE): retinal pigment epithelial cells
(RXRs): retinoic X receptor
(VEGF): vascular endothelial growth factor

